# Quantitative Ultrasound Analysis of Gas Embolic Disease in Bycaught Sea Turtles

**DOI:** 10.64898/2026.07.27.740119

**Authors:** Katherine Eltz, Jose-Luis Crespo, Arian Azarang, Emma Pla-González, Daniel García-Párraga, Andreas Fahlman, Virginie Papadopoulou

**Affiliations:** Department of Radiology, The University of North Carolina at Chapel Hill, North Carolina, USA; Lampe Joint Department of Biomedical Engineering, The University of North Carolina at Chapel Hill & North Carolina State University, North Carolina, USA; Fundación Oceanogràfic de la Comunitat Valenciana, Valencia, Spain; Linköping University, IFM, Linköping, Sweden

**Keywords:** Gas embolism, Texture, Decompression, Decompression sickness, Imaging biomarker, Image analysis, Venous gas emboli, Ultrasound imaging

## Abstract

**Objectives:** Bycatch-related decompression after forced submersion can result in severe gas embolic disease in sea turtles. This work used qualitative and quantitative ultrasound analyses, including gas grading, brightness analysis, and texture feature extraction, to investigate organ-specific gas burden and temporal evolution in the hearts, kidneys, and livers of bycaught sea turtles.

**Materials and Methods:** Ultrasound imaging of the hearts, kidneys, and livers of bycaught turtles was performed as part of veterinary evaluation either onboard fishing vessels immediately after surfacing (boat group, n=47) or after longer periods at shore-based facilities (shore group, n=30). Gas burden in each ultrasound scan was graded on an ordinal scale from 0 (no gas) to 5 (gas completely shadowing organ anatomy). Temporal differences in gas burden were compared between the shore and boat groups. Quantitative brightness and texture features were extracted from all organs, including contrast, correlation, homogeneity, and energy from liver and kidney data. A multivariate logistic regression model with leave-one-out cross-validation was conducted, with shore versus boat as a binary outcome (surrogate of post-surfacing decompression state) and ultrasound texture metrics as independent variables.

**Results:** Median grades from the first ultrasound scan were significantly higher in the boat group than in the shore group for the liver, kidney, and heart (3, 3, and 3 vs 1, 1, and 0, respectively). This pattern coincided with a difference in the mean duration until the first scan which was conducted being 54 min for the onboard studies vs 330 minutes in the shore group. Mean pixel brightness within cardiac and liver regions of interest increased with rising bubble grade before decreasing at the highest grades, consistent with acoustic shadowing at severe gas burden. Texture features demonstrated significant organ-specific changes with increasing gas burden, and the regression models achieved areas under the receiver operating characteristic curve of 0.92 for liver texture features and 0.83 for kidney texture features.

**Conclusions:** These findings demonstrate organ-specific differences in gas evolution over time. Quantitative ultrasound features were associated with gas burden and post-surfacing interval in bycaught sea turtles. These findings support the feasibility of quantitative ultrasound biomarkers for assessment of decompression-related gas burden.

## INTRODUCTION

Decompression sickness (DCS) results from reductions in ambient pressure leading to inert gas supersaturation and bubble formation in blood and tissues.^1,2^ In human diving operations, DCS remains a major limitation in diving operations and hyperbaric environments^3,4^, and symptoms range from mild musculoskeletal pain to severe neurologic injury and death.^3,4^ A number of risk factors have been identified, including dive depth and duration. It is also well established that larger animals are at greater risk than smaller animals undertaking the same dive profile, likely because body size influences gas uptake and elimination dynamics through its effects on blood flow and tissue gas exchange.^5^ Despite this, DCS susceptibility remains difficult to predict at the individual level. Venous gas emboli (VGE) detected by ultrasound are therefore widely used as surrogate markers of decompression stress in research settings. However, while increasing VGE grade is associated with elevated DCS risk at the population level, VGE has limited positive predictive value for clinical DCS, and asymptomatic dives with detectable VGE are common even after controlled exposures.^6^ Importantly, even operational military and experimental diving protocols are typically designed to maintain symptomatic DCS rates below 2%, inherently limiting acquisition of human datasets with severe systemic gas burden.^7^

Ultrasound-based VGE assessment is most commonly performed using Doppler ultrasound or transthoracic echocardiography, with bubble burden graded by trained human operators using ordinal systems such as the Spencer or Eftedal-Brubakk scales (Table S1, Supplemental Digital Content 1). This approach remains central to decompression research because increasing VGE severity is associated with higher DCS risk.^8^ Nevertheless, current ultrasound assessment remains predominantly qualitative and observer dependent, with known variability between individuals performing the scoring, devices, and acquisition techniques. Moreover, most human decompression datasets are derived from controlled diving studies involving relatively mild-to-moderate decompression stress and solely intravascular gas burden. Both Doppler ultrasound and echocardiographic bubble grading are fundamentally limited to detection of moving intravascular gas.^9^ As a result, imaging characteristics of static or tissue-associated gas burden remain poorly characterized in human decompression studies. Ethical and operational constraints inherently limit acquisition of human datasets containing severe systemic gas embolic disease with extensive tissue gas accumulation. Consequently, the imaging characteristics and temporal evolution of high gas burden states remain incompletely understood.

Current understanding of decompression pathophysiology suggests that gas burden evolves differently across tissues and organ systems depending on local perfusion, tissue gas kinetics, and supersaturation profiles.^5^ VGE may appear within minutes after surfacing and persist for hours, while slower tissues remain supersaturated longer and may exhibit prolonged or delayed gas accumulation depending on the depth-time exposure profile. Organ-specific injury patterns have been described in the spinal cord, lungs, skin, kidneys, liver, and inner ear, highlighting the heterogeneous nature of decompression-related pathology.^1^ However, characterization of these processes has relied largely on qualitative ultrasound grading, indirect physiologic metrics, or provocative experimental animal models that result in high gas burden and DCS rates. Quantitative imaging biomarkers capable of objectively assessing gas burden and temporal evolution remain limited.

Breath-hold diving marine animals provide a unique opportunity to study naturally occurring severe gas embolic disease. Under normal conditions, sea turtles possess physiologic adaptations that limit nitrogen uptake during diving, including, incomplete interventricular septum, increased smooth muscle in the sinus venosus to regulate atrial and ventricular filling, and sphincters in the pulmonary arteries that helps to regulate pulmonary resistance and blood flow.^10,11^ However, extended forced submersion and subsequent ascent during bycatch can disrupt these protective mechanisms and lead to systemic gas emboli formation.^12^ Human activities are the primary cause of sea turtle endangerment, with 5 of the 7 sea turtle species currently classified as threatened or endangered, underscoring the importance of understanding the long-term effects of bycatch events on affected animals.^13^ Previous studies have shown that paralysis, lethargy, and eventual death can occur after surfacing, and therefore release following bycatch-related gas embolism can result in unrecorded animal death.^12,14,15^ Unlike controlled human diving studies, these animals may present with extensive systemic gas accumulation in both vascular and tissue compartments, offering a rare naturally occurring model of severe decompression injury.

Prior work in sea turtles has primarily focused on the presence of gas emboli, clinical management, and associated mortality following bycatch. Previous studies have also demonstrated associations between fishing depth, body size, and gas embolism severity in sea turtles, with gas visualized using radiography and ultrasound in the heart, liver, kidneys, and major vessels.^16–18^ Such studies have suggested that gas resolves at different rates across organ systems, with cardiac intravascular gas decreasing more rapidly than gas within renal and peripheral vascular structures, and radiographic evidence of gas emboli reported in 42.4% of 482 bycaught turtles.^17^ However, organ-specific temporal evolution of gas burden has not been quantitatively characterized with ultrasound. In addition, most existing assessments remain dependent on subjective visual interpretation, analogous to limitations encountered in ultrasound grading following human decompression. Quantitative ultrasound features capable of capturing embolic burden and spatial heterogeneity may therefore improve characterization of decompression-related pathology across a broader spectrum of disease severity.

In this study, we used ultrasound imaging to characterize temporal changes in gas burden in bycaught sea turtles following surfacing. In addition to qualitative ultrasound grading, we applied quantitative image analysis methods including brightness analysis and texture feature extraction to ultrasound images of the heart, liver, and kidneys. Because gas strongly reflects ultrasound, image brightness may reflect embolic burden, whereas texture features may capture spatial heterogeneity while reducing sensitivity to acquisition variability. By leveraging a naturally occurring dataset with high systemic gas burden, we evaluated quantitative ultrasound features that are difficult to study in human DCS cohorts. These findings can help development of quantitative ultrasound biomarkers for gas embolic disease and inform future ultrasound-based approaches in decompression research.

## METHODS

### Imaging Methods

Ultrasound imaging of the hearts, kidneys and livers of loggerhead (n = 77) was previously performed in the field as part of veterinary assessment in bycaught wild animals, as well as in animals presenting on shore and brought to veterinary centers for assessment and treatment. The study sample was categorized into two distinct groups: those imaged immediately on fishing vessels post-surfacing (boat group, n = 47) and those imaged after transport to shore-based facilities (shore group, n = 30).

Ultrasound data were collected between 2018 and 2023 in Italy, Spain, and Brazil, using a GE Logiq V ultrasound scanner and curvilinear probe for cardiac, hepatic, and renal imaging, with additional linear probe kidney acquisitions. The sea turtles underwent ultrasonography through the acoustic windows necessary to access the heart, liver, and kidney, using the ventral and lateral cervical regions, the axillary access, and the cranial region of the inguinal acoustic window, respectively. Ultrasound images were collected at frequencies between 4 and 10 MHz, gain values between 42 and 67, and depths ranging from 3 to 30 centimeters. Intra-subject settings were typically unchanged from one timepoint to the next; the setting ranges stated above account for turtles of varying sizes and target organs of different depths. A schematic showing the data collection timeline is provided in Figure 1.

**Figure 1:**
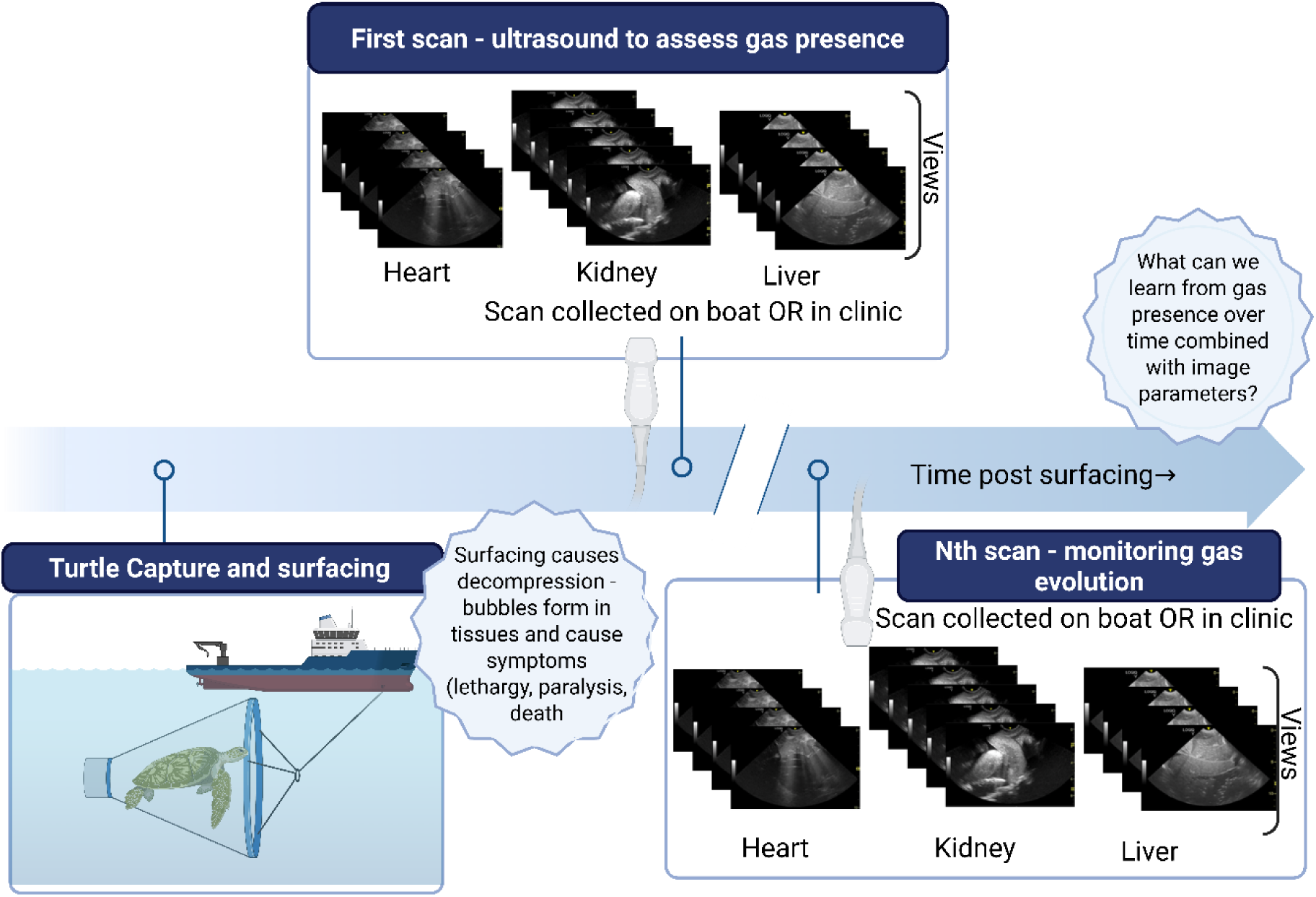
Schematic of the data collection timeline following bycatch. Ultrasound imaging of the hearts, kidneys, and livers was performed either onboard the vessel or at a veterinary facility, corresponding to the two datasets analyzed. Multiple views were acquired for each organ at each time point.

### Quantitative data analysis

Gas quantity in each ultrasound scan was graded on an ordinal from 0 (no gas) to 5 (gas completely shadowing organ anatomy) by a trained veterinarian with experience in turtle ultrasonography (JLC). During the grading process, scans were examined for features that indicate high levels of gas, such as bubbles, B-lines, and shadowing. The grading was based on the presence or absence of bubbles in the bloodstream and tissues, the subjective number of both moving and stationary bubbles, and the generation of artefacts such as comet tails. In the most severe cases, the examination was prevented by the large amount of gas present. Grading criteria are summarized in Table 1.

**Table 1:** Grading scale for assessing gas presence in turtle ultrasound scans. Grades are based on the amount of gas and reflect the severity of gas embolic pathology based on radiologic findings, as assessed by trained veterinarians.

|  |  |
| --- | --- |
| 0 | No visible gas |
| 1 | Slight presence of bubbles circulating |
| 2 | Constant presence of bubbles in blood flow |
| 3 | Higher presence of bubbles circulating, B-line artifacts visible |
| 4 | Massive presence of bubbles in blood flow |
| 5 | Clear images cannot be obtained due to the large amount of gas |

The time of surfacing and the interval between surfacing and ultrasound acquisition were retrieved for each turtle. Data were organized into two groups based on ultrasound collection location (“shore” and “boat”). After confirming temporal differences between groups, the initial recorded grade for each organ was analyzed to assess immediate post-surfacing gas burden. To evaluate gas progression over the entire observation period, all recorded grades were aggregated into 100-minute intervals (Figure S1, Supplemental Digital Content 2).

### Development of graphical user interface pipeline

The dataset included multiple timepoints per organ, with each timepoint containing several imaging views. To facilitate standardized handling of the ultrasound dataset, a Python-based graphical user interface (GUI; Tkinter, available at ^19^) was developed. The GUI enabled selection of representative views, removal of duplicate acquisitions, temporal cropping to exclude motion artifacts, and manual delineation of anatomical regions of interest (ROI) for quantitative analysis. These features are detailed in Table 2, and the GUI workflow is shown in Figure 2.

**Figure 2:**
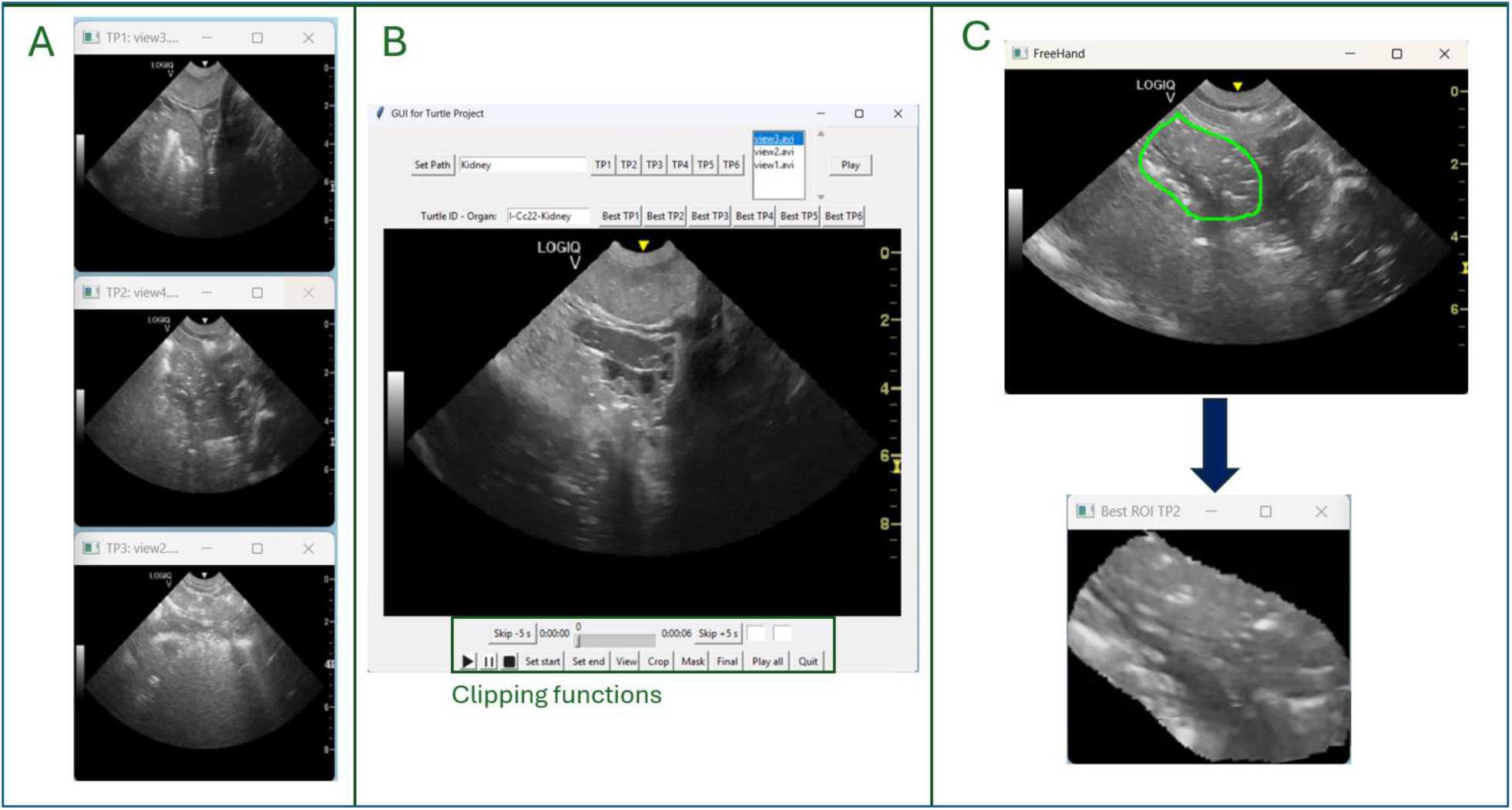
The graphical user interface used in analysis. A) Users can view multiple timepoints from one dataset at once to find consistent videos. B) After selecting the best video from each timepoint, clipping features can be used to remove motion artifacts. C) Users can draw regions of interest around output videos from step B, repeating for each timepoint, for use in future analysis.

**Table 2:** Overview of graphical user interface functions to address dataset variability.

| Sources of variation in the data | How the variations are mitigated through the use of the graphical user interface (GUI) |
| --- | --- |
| Multiple videos per timepoint | Simultaneous visualization of all videos from a timepoint to identify the most consistent acquisitions |
| Motion artifacts | Temporal cropping to stationary video segments (typically 1–3 seconds) |
| Unnecessary anatomy included in scans | Manual ROI delineation around target anatomy |
| Differences in frequency, gain, and depth | Not directly corrected within the GUI |

Because acquisition parameters including probe position, gain, imaging depth, and frequency varied owing to the field-based nature of data collection, the GUI was designed to reduce non-gas-related sources of imaging variability. Residual acquisition variability was addressed in subsequent quantitative image analysis steps.

### Pixel Brightness Analysis

ROIs were defined differently for each organ. In higher-grade cardiac and liver scans, anatomical structures were frequently obscured by gas-related shadowing. Therefore, ROIs for cardiac and liver scans were standardized as the central column of pixels in the upper half of the imaging cone. For kidney scans, visible vessels were selected as the ROI when identifiable. Mean pixel brightness was calculated within each ROI using MATLAB.

### Ultrasound Image Texture Analysis

Haralick texture features, which characterize spatial brightness relationships beyond global image intensity, were used as an additional quantitative metric for gas characterization across variable ultrasound acquisition settings.^20^ Four texture features: contrast, correlation, energy, and homogeneity, were evaluated.

Texture features were derived from gray-level co-occurrence matrices (GLCMs), which quantify the frequency of neighboring pixel intensity combinations within an image. GLCMs were computed in MATLAB using parameter ranges selected from prior ultrasound texture-analysis studies^21–24^, including step sizes of 1–5 pixels, step angles of 0°, 45°, 90°, and 135°, and gray level bin counts of 16 or 32. The equations used to compute each texture feature are shown below, where *i* and *j* represent matrix indices corresponding to gray-level intensity pairings, *p(i,j)* correspond to the elements of the normalized GLCM, and *σ_i_* and *σ_j_* correspond to the standard deviations of the GLCM data.

Contrast :

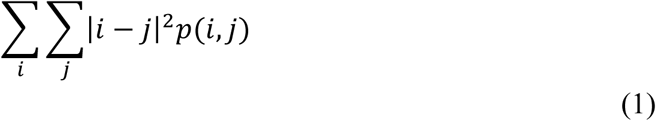

Correlation:

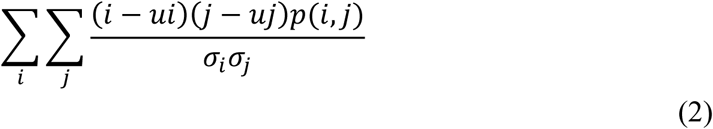

Energy:

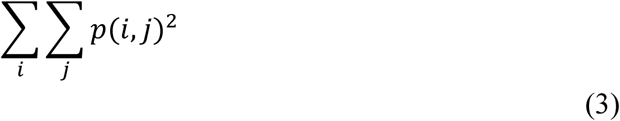

Homogeneity:

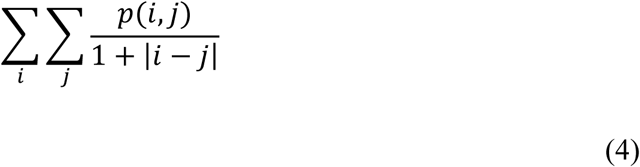

Texture analysis was performed on 50 x 60-pixel rectangular ROIs within kidney and liver tissue free from major motion artifacts. Texture features were averaged across all frames within each video. Resulting values were subsequently grouped by grade to evaluate differences in texture parameters. Representative ROIs used for kidney and liver texture analysis are shown in Figure S2, Supplemental Digital Content 3.

### Statistical analyses

All statistical analyses were performed using GraphPad Prism (Version 10.0, GraphPad Software, Inc. San Diego, CA, USA.) or MATLAB (R2025b. The MathWorks Inc., Natick, MA, 2025) and a threshold of p < 0.05 was set *a priori* and considered statistically significant.

The time delay between surfacing and initial ultrasound acquisition for the shore and boat groups was analyzed using a Mann-Whitney test. Comparisons of median initial organ grades between shore and boat groups were performed using Mann-Whitney testing. Differences in brightness across organs were assessed using a Kruskal-Wallis test, followed by Dunn’s post hoc analysis (p < 0.05), to compare grade 0 with each subsequent grade. Texture parameters were similarly analyzed using a Kruskal–Wallis test with grade 0 as reference (α < 0.05). Linear regression analysis was performed to determine whether parameter slopes differed significantly from zero.

Multivariabte logistic regressions with leave-one-out cross-validation were performed using texture features as predictors to assess the ability of liver and kidney texture parameters to classify scan location (shore versus boat) as a proxy for time delay. Model discrimination was assessed using receiver operating characteristic (ROC) analysis, with areas under the curve (AUCs) and corresponding 95% confidence intervals calculated from leave-one-out cross-validated predictions. Correlation and collinearity analyses were additionally performed to evaluate redundancy and dependency among texture features. Finally, principal component analysis (PCA) was used to assess clustering of ultrasound datasets from shore and boat acquisitions based on texture-feature distributions.

## RESULTS

### Timing differences and gas burden evolution between shore and boat groups

A comparison of the interval between surfacing and first ultrasound acquisition for turtles in the boat and shore groups is shown in Figure 3A. The first imaging timepoint occurred significantly later in the shore group (median = 330 minutes, range = 90 - 1440 minutes, n = 30, Fig. 3) compared to the boat group (median = 54 minutes, range: 6 - 1307 minutes, n = 49) Mann-Whitney test).

**Figure 3:**
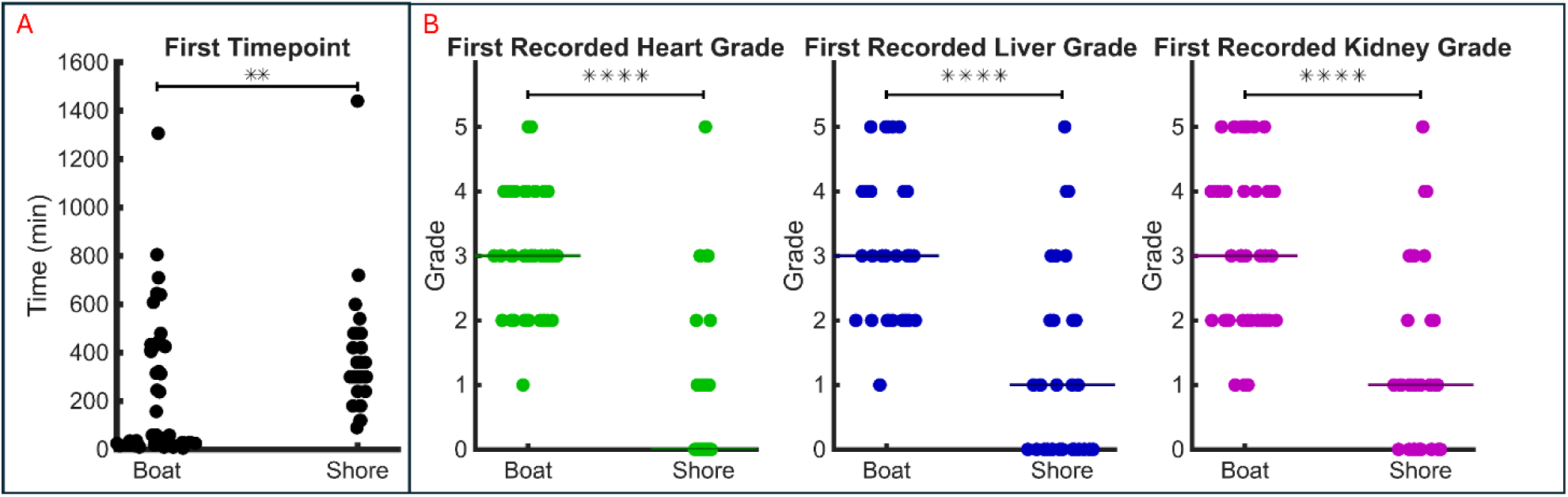
A) Average times between surfacing and first ultrasound acquisition for all animals separated by shore and boat groups. B) Distribution of initial ultrasound grades separated by shore and boat groups.

Initial gas grades of animals captured only in trawling nets were subsequently compared by organ (Figure 3B). The median first grade in the heart, liver, and kidney, for the shore group was 0, 1, and 1, respectively, compared with 3, 3, and 3 in the boat group. Differences between groups were significant for all organs (Mann–Whitney test). Sensitivity analyses including all animals, regardless of capture method or surface interval availability, are presented in Figure S3 (Supplemental Digital Content 4) and demonstrated the same significant differences between the boat and shore groups.

Temporal evolution of gas burden is shown in Figure S1, Supplemental Digital Content 2, in which grades were grouped into 100-minute intervals post-surfacing. Notably, grade 5 scans, indicative of extensive gas accumulation, persisted longer in the kidneys and liver than in the heart.

### Qualitative imaging findings

Representative examples of temporal gas evolution within the kidney, heart, and liver from individual animals are shown in Figure 4. Increased gas burden was visually associated with increased echogenicity within vascular structures and progressive acoustic shadowing at the highest burdens. In kidney scans, increasing grade was associated with increasing B-line artifacts and hyperechoic intravascular signals (Figure 4A). In some cardiac scans, higher grades demonstrated progressively greater intravascular hyperechoic bubble burden while preservation of chamber anatomy remained visible in some grade 4 acquisitions (Figure 4B). In some liver and cardiac scans, increasing gas burden produced progressive acoustic shadowing, resulting in partial or complete obscuration of deeper anatomy at higher grades (Figure 4C).

**Figure 4:**
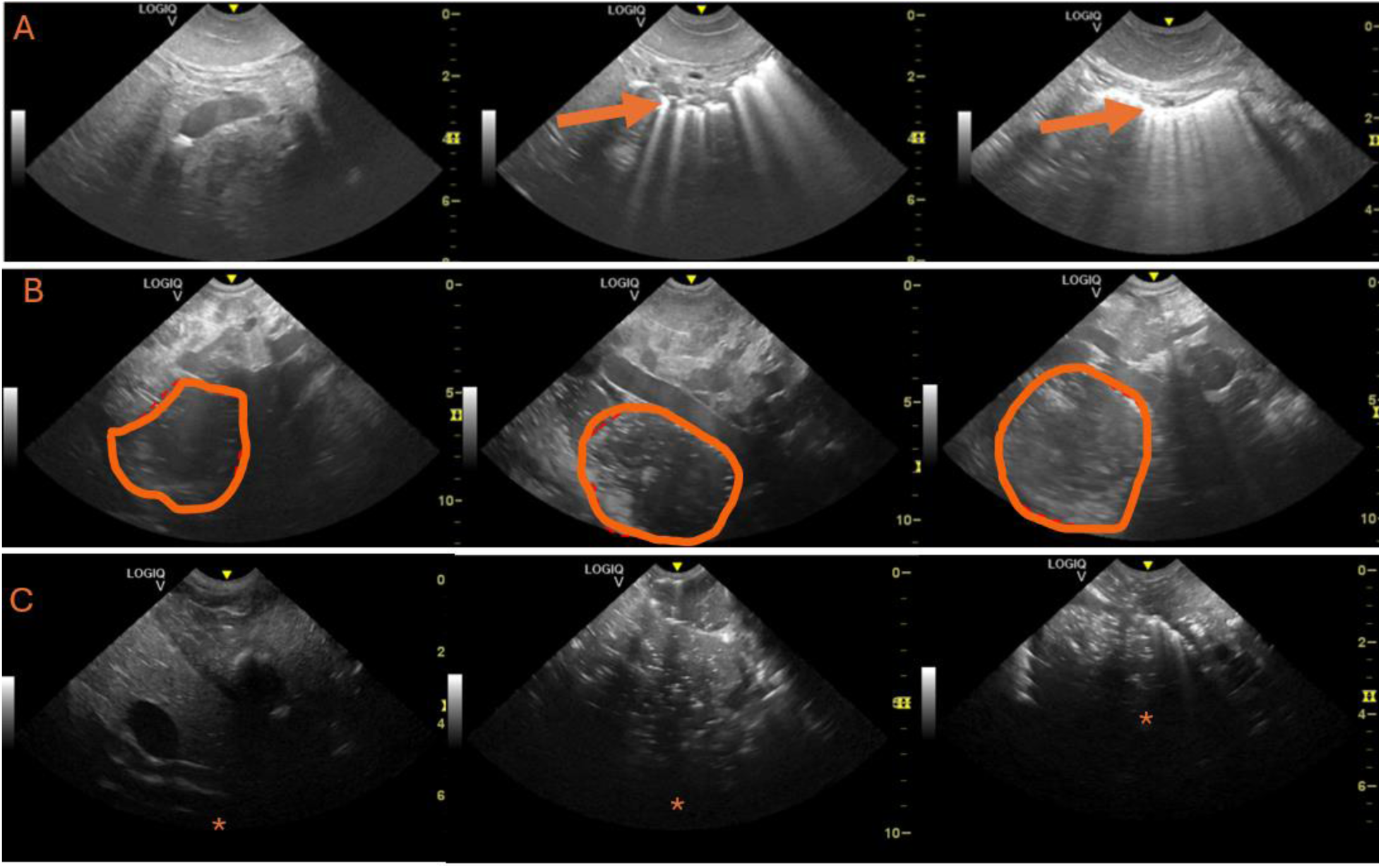
Representative ultrasound frames demonstrating qualitative imaging changes associated with increasing gas burden. A) Kidney grades 2, 3, and 4, acquired at 59, 115, and 163 minutes post surfacing, respectively. The appearance of B-lines (indicated with orange arrows) increases with grade, indicating the presence of higher amounts of bubbles. B) Heart grades 2, 3, and 4, acquired at 19, 59, and 79 minutes post surfacing, respectively. Orange lines encircle one heart chamber, with higher amounts of bright spots (bubbles) as grade increases. C) Liver grades 2, 4 and 5, acquired at 15, 40, and 95 minutes post surfacing, respectively. Increased shadowing is apparent at higher grades, as a darkened lower portion of each image (indicated by the orange star depth). For each separate organ timeseries shown in A-C, the depicted frames are all taken from the same animal over time. Each organ time series in A-C was obtained from a single animal.

### Brightness metrics

Brightness analysis results from each organ are shown in Figure 5. In cardiac and liver datasets, mean ROI brightness generally increased with increasing grade before decreasing at the highest grades. This pattern was consistent with increasing gas reflectivity followed by acoustic shadowing at severe gas burden, where overwhelming gas accumulation acted as a specular reflector and impeded acoustic penetration. In kidney vessels, increasing grades were associated with a broader distribution and greater variability of brightness values without a corresponding monotonic increase in median brightness.

**Figure 5:**
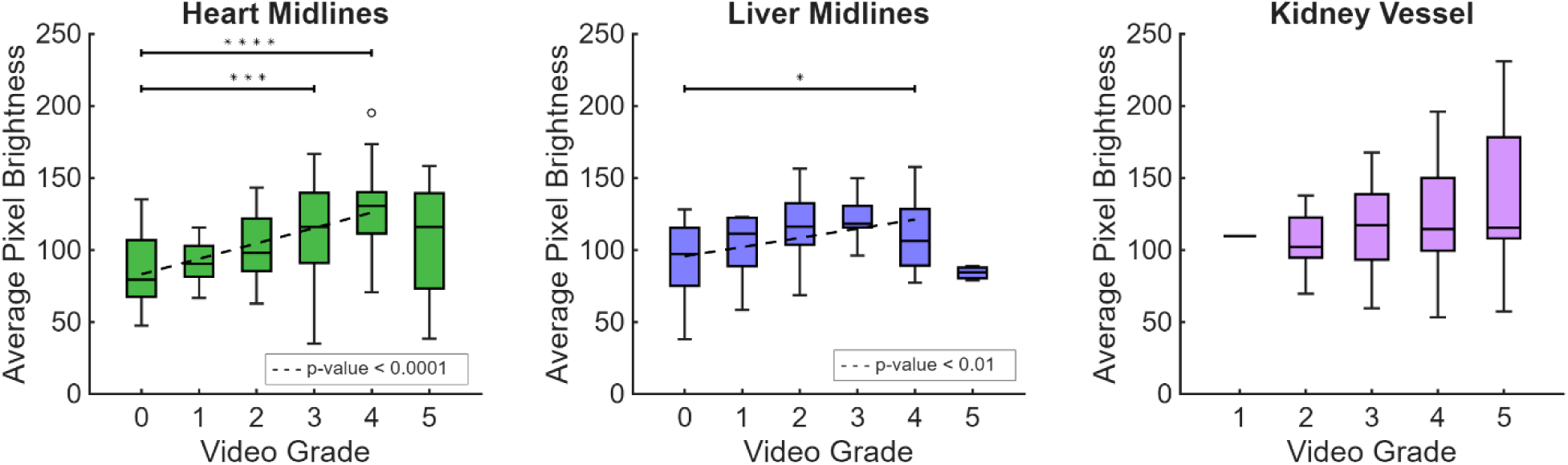
Mean pixel brightness values within organ-specific regions of interest (ROI) grouped by grade.

### Texture metrics

Using five step sizes, four step directions (plus the mean angle), and either 16 or 32 gray levels, a total of 50 gray-level co-occurrence matrices were generated per image. Four texture features (contrast, correlation, energy, and homogeneity) were calculated from each matrix. Only the most robust parameter combinations are reported here, corresponding to 32 gray levels, a step size of 2, and averaged angular analysis. Increasing gas grade was associated with significant increases in contrast and significant decreases in homogeneity and energy in both liver and kidney datasets (Figure 6).

**Figure 6:**
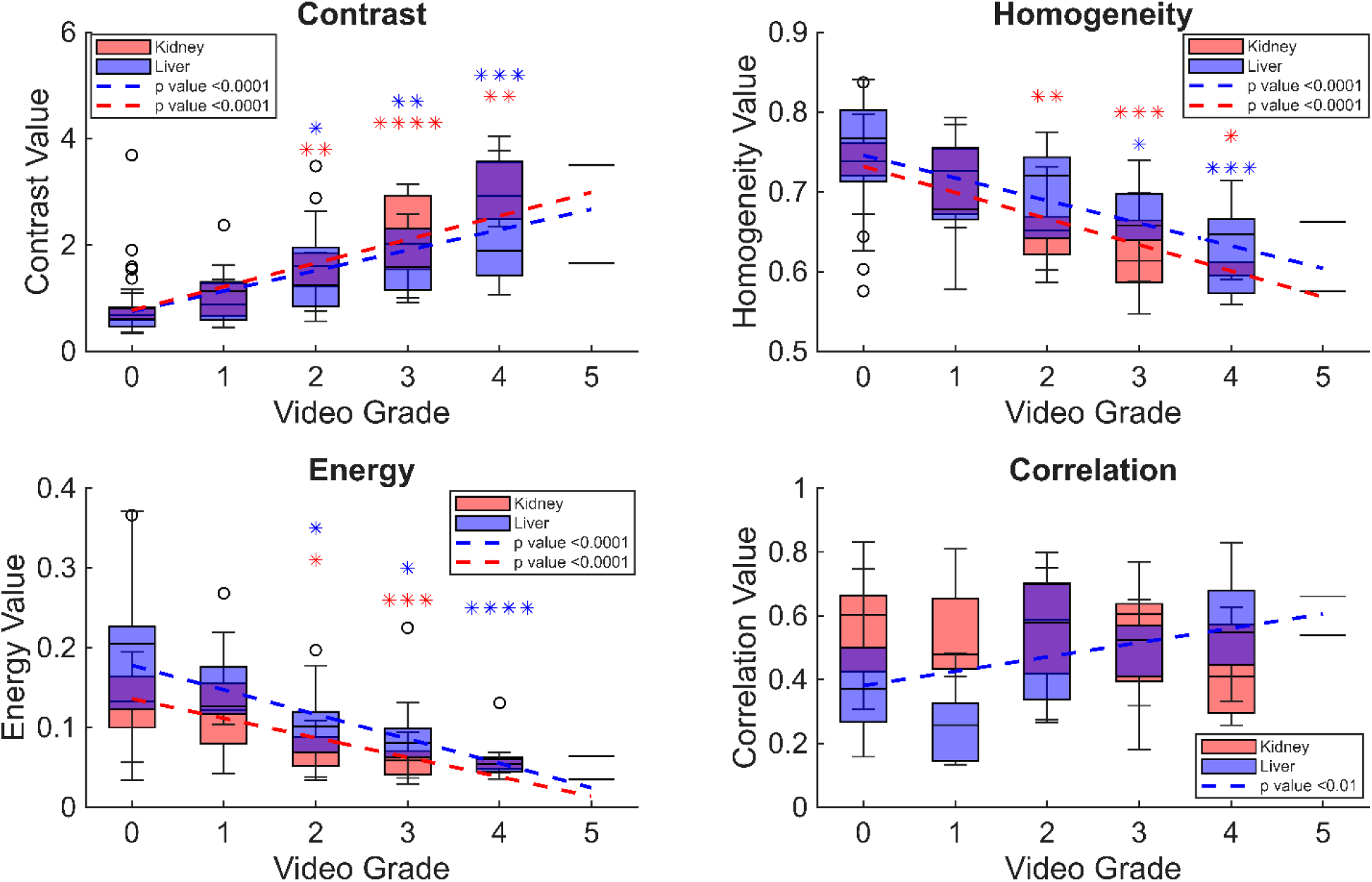
Texture-feature analysis results for liver (blue) and kidney (red) datasets. Significant increasing trends with grade were observed for contrast, whereas significant decreasing trends were observed for energy and homogeneity. Examples of grade 5 were n = 1 for every case, so that data point was excluded from this analysis. Texture feature values per grade significantly different from grade 0 are shown with * for liver (blue) and kidney (red) respectively (Kruskal-Wallis with Dunn’s post-hoc test, p < 0.05).

### Logistic Regression and multivariate analysis

Multivariate logistic regression with leave-one-out cross-validation was performed using texture-feature combinations to predict scan location (shore versus boat) in liver and kidney datasets. Only statistically significant texture features were included in regression analysis, consisting of all four texture features for liver data and contrast and energy for kidney data. Cross-validated models demonstrated strong discriminative performance, with AUCs of 0.92 (95% CI: 0.72 – 0.99) for liver texture features and 0.83 (95% CI: 0.68 – 0.93) for kidney texture features. Corresponding ROC curves are shown in Figure 7.

**Figure 7:**
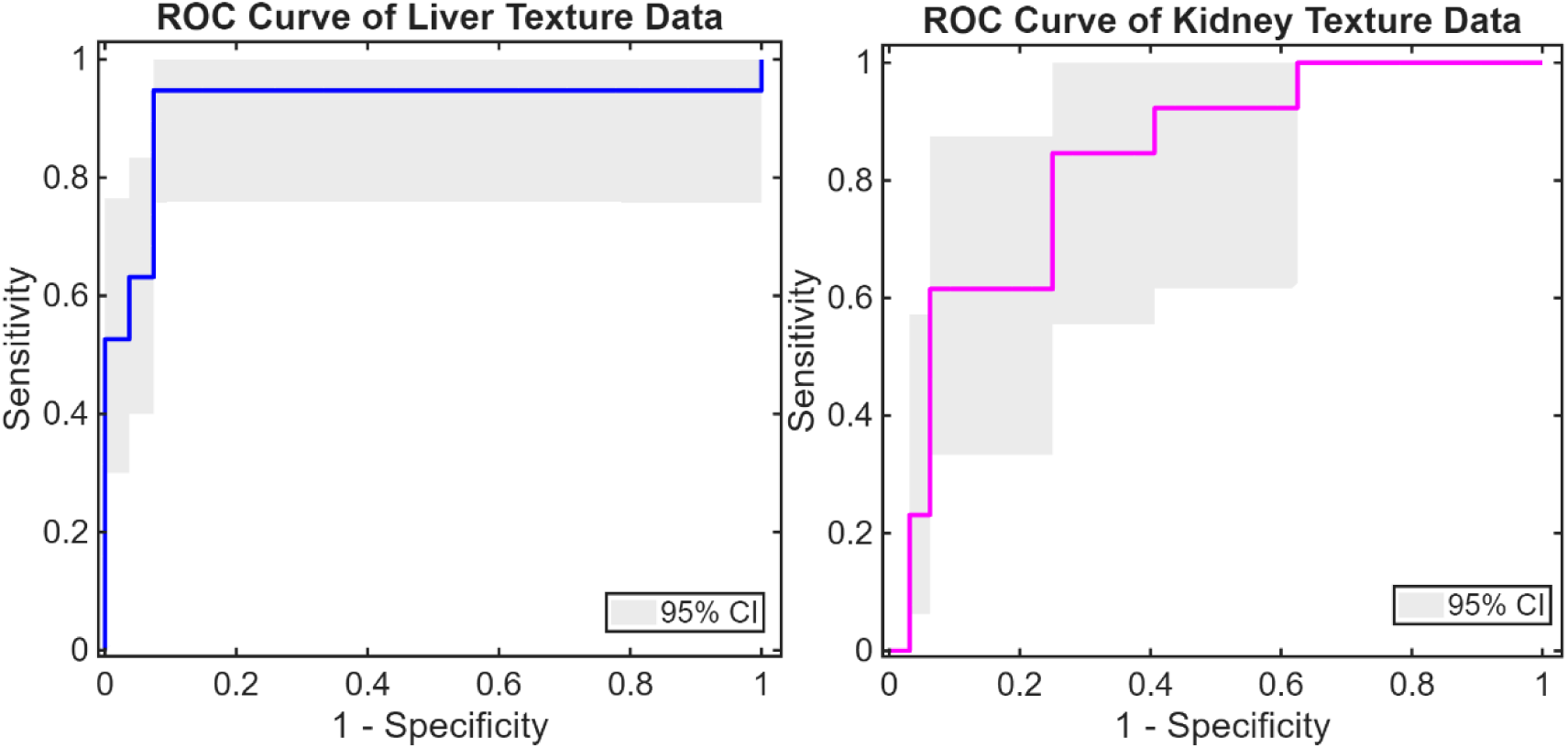
Receiver operating characteristic (ROC) curves for shore-versus-boat classification using liver and kidney texture features. AUCs are 0.92 (95% CI: 0.72 – 0.99) for liver texture features and 0.83 (95% CI: 0.68 – 0.93) for kidney texture features.

Leave-one-out cross-validation was performed such that each prediction was generated using a model trained without the corresponding test datapoint. Logistic regression models generated using the full dataset were additionally evaluated to assess feature correlation and collinearity. Covariance matrices are shown in Table S2, Supplemental Digital Content 5. In both liver and kidney datasets, energy and homogeneity demonstrated the lowest redundancy, whereas contrast and energy showed the greatest redundancy. Variance inflation factor analysis did not demonstrate problematic multicollinearity among texture features.

Finally, principal component analysis was also performed using statistically significant texture features to evaluate clustering of shore and boat datasets. Results demonstrated partial clustering between groups (Figure S4, Supplemental Digital Content 6).

### DISCUSSION

This study demonstrates that quantitative features from ultrasound imaging can characterize temporal and organ-specific differences in gas burden in bycaught sea turtles following surfacing. Gas accumulation evolved differently across organs, with persistent high-grade gas remaining longer in the kidneys and liver than in the heart. In addition, quantitative brightness and texture-derived image features correlated with qualitative gas severity despite heterogeneous field-acquisition conditions. These findings support the feasibility of quantitative ultrasound biomarkers for assessing decompression-related gas burden beyond conventional qualitative grading of intravascular bubbles. ^9,25^

The observed differences in gas persistence between organs are consistent with prior work describing heterogeneous gas clearance dynamics in bycaught sea turtles. Franchini et al. previously demonstrated faster radiographic gas disappearance from the heart and major vessels compared with renal and iliac vasculature.^17^ In that study, full-body radiographs were obtained every 48 hours until either gas resolution or death, and the earliest signs of gas disappearance consistently occurred within the heart, venous sinus, and great vessels, whereas renal and iliac vessels were among the last structures to normalize. Similarly, the present study demonstrated lower cardiac grades at delayed shore-based imaging timepoints and prolonged severe gas burden in the kidneys and liver. These findings support the concept that decompression-related gas evolution differs substantially across organ systems, likely reflecting differences in local perfusion, vascular architecture, tissue gas kinetics, and tissue supersaturation profiles^26^.

Importantly, current decompression ultrasound methods primarily assess moving intravascular bubbles within the bloodstream using Doppler ultrasound or transthoracic echocardiography. Both modalities fundamentally rely on motion-based detection of highly reflective gas interfaces and therefore incompletely characterize static or tissue-associated gas accumulation.^9^ Prior decompression ultrasound studies have repeatedly shown that venous gas emboli (VGE) are common following asymptomatic dives and only imperfectly correlate with clinical decompression sickness (DCS).^4,8^ Furthermore, experimental diving protocols are generally designed to maintain low clinical DCS rates, inherently limiting access to human datasets with severe systemic gas burden and extensive tissue gas accumulation. However, VGE amounts as detected by ultrasound have been correlated to mortality and to systemic gas burden in post-mortem evaluations in a small mammal model.^27^ That study emphasizes the importance of ultrasound in detecting VGE and differentiating high gas states in afflicted animals, as the imaging characteristics of high-burden decompression states remain poorly understood. The naturally occurring pathology observed in bycaught sea turtles therefore provides a unique opportunity to investigate severe decompression-related gas accumulation involving both vascular and tissue compartments.

The prolonged renal gas burden observed in this study may be particularly relevant mechanistically. Prior turtle studies have suggested that vascular gas may persist longer within renal and iliac systems than within the heart ^17^, possibly reflecting slower washout kinetics or retention within peripheral vascular beds. Influence of anatomical reptilian portal renal system present in sea turtles draining an important portion of the venous circulation from the caudal body region through kidneys before reaching the postcava most probably favors gas presence in this region.^28,29^ In the present dataset, kidneys consistently demonstrated persistent higher-grade findings relative to cardiac imaging at delayed imaging timepoints. These observations support the hypothesis that renal ultrasound may provide a particularly sensitive imaging target for prolonged decompression-related gas burden in animals with a renal portal vascular system.^28^ More broadly, these findings suggest that decompression-related ultrasound changes may reflect not only intravascular bubbles but also evolving tissue–gas interactions occurring over time following surfacing.

Brightness analysis demonstrated organ-dependent relationships between quantitative intensity measurements and gas severity. In the heart and liver, brightness initially increased with gas grade and subsequently decreased at severe grades because overwhelming gas accumulation produced extensive acoustic shadowing and limited ultrasound penetration. This behavior is consistent with the large acoustic impedance mismatch between gas and surrounding tissue, which causes strong reflection and attenuation of the incident ultrasound wave. Similar shadowing phenomena have previously been reported in decompression and contrast-enhanced ultrasound literature, where high microbubble concentrations attenuate deeper acoustic signal transmission. ^25,30^ Recent work in a porcine decompression model similarly demonstrated reduced ultrasound brightness in lymphatic imaging regions in association with severe DCS and gas accumulation.^25^ In contrast, kidney vessel ROIs in the present study demonstrated greater variability and broader brightness distributions without a monotonic increase in median brightness. Because these ROIs primarily sampled more superficial vascular structures rather than deeper shadowed tissue, the measured signal likely reflected heterogeneous intravascular gas distributions (both spatially and temporally) rather than secondary attenuation effects. Together, these findings suggest that brightness measurements may provide a useful quantitative surrogate for gas burden in mild-to-moderate decompression states while becoming increasingly influenced by acoustic saturation effects at severe grades.

A major challenge in decompression ultrasound research is the substantial variability introduced by acquisition parameters including gain, imaging depth, frequency, focal position, and proprietary system processing.^9^ These factors complicate quantitative comparisons across datasets and may obscure subtle decompression-related image changes. In the present study, acquisition variability was partially mitigated through standardized ROI selection, temporal clipping, and texture-feature analysis designed to capture local spatial relationships rather than global image intensity alone. The use of a graphical user interface pipeline additionally improved consistency in scan selection and artifact exclusion across longitudinal imaging datasets. Other post hoc standardization approaches have recently been applied in decompression ultrasound investigations of lymphatic bubbles, where phantom-based brightness correction substantially reduced variability introduced by nonuniform acquisition settings.^25^

Texture analysis provided additional quantitative characterization of decompression-related ultrasound changes and may offer advantages over brightness analysis under heterogeneous field conditions. Haralick texture features evaluate spatial relationships between neighboring pixel intensities rather than isolated pixel amplitude values, potentially reducing sensitivity to acquisition variability. Increasing gas burden was associated with increased contrast and decreased homogeneity and energy in both liver and kidney datasets, indicating progressively heterogeneous spatial intensity distributions as gas burden increased. These findings likely reflect altered ultrasound speckle patterns and scattering behavior caused by gas– tissue interactions rather than exclusively visualization of discrete bubbles. Importantly, texture alterations remained detectable even in scans with substantial shadowing and reduced anatomical conspicuity. This observation suggests that quantitative texture features may capture imaging signatures of severe gas accumulation that are difficult to evaluate qualitatively using conventional grading approaches alone.

The multivariate logistic regression analyses further support the potential utility of quantitative image features for characterizing decompression-related imaging states. Cross-validated models demonstrated strong discriminative performance for classification of shore versus boat imaging groups, which in this dataset directly reflected post-surfacing imaging delay and gas burden evolution. These findings suggest that texture features encode temporally associated decompression-related image changes linked to gas burden evolution over time. Liver texture models demonstrated greater performance than kidney models, possibly reflecting more pronounced shadowing and heterogeneity associated with severe hepatic gas accumulation. The relationship between texture features and temporal decompression state additionally suggests potential utility for estimating post-surfacing gas evolution in situations where limited exposure history is available at presentation.

Several limitations should be acknowledged. First, this was a retrospective analysis of field-acquired ultrasound data originally obtained for veterinary management rather than standardized quantitative imaging research. Consequently, although using the same equipment and team of professionals, acquisition parameters including gain, frequency, imaging depth, probe positioning, and time sampling varied across examinations. Second, severe gas burden frequently obscured anatomical structures because of acoustic shadowing, particularly in grades 4 and 5. Third, only selected quantitative image features were evaluated, and additional approaches including vessel-flow analysis, attenuation mapping, automated B-line quantification, or nonlinear ultrasound methods may provide complementary information. Recent decompression ultrasound developments have emphasized the importance of nonlinear and contrast-specific imaging approaches for evaluation of microbubble behavior and stationary tissue-associated gas.^9^ The logistic-regression analyses were internally validated using leave-one-out cross-validation without external validation in an independent dataset. Larger datasets with more standardized acquisition protocols across individual turtles would further strengthen the generalizability of these findings. Finally, this reptilian model contains varying anatomy and physiology when compared to mammal models, so gas tissue dynamics investigated here may not be directly applicable to all taxa.

Despite these limitations, the present work demonstrates the feasibility of quantitative ultrasound assessment of severe decompression-related gas burden in a naturally occurring animal model. By combining longitudinal imaging, organ-specific assessment, and quantitative image-feature extraction, this study extends decompression ultrasound analysis beyond conventional qualitative grading of moving intravascular bubbles. These findings provide a framework for future investigation of quantitative ultrasound biomarkers of decompression-related gas accumulation in both veterinary and human decompression research.

## CONCLUSION

This study demonstrates that quantitative ultrasound metrics, including brightness and texture-based features, can characterize temporal and organ-specific differences in gas burden in bycaught sea turtles following surfacing after forced submersion. Gas persistence differed across organs, with prolonged severe gas burden observed in renal and hepatic imaging compared with cardiac imaging. Quantitative image features correlated with qualitative gas severity and remained informative despite heterogeneous acquisition conditions and extensive acoustic shadowing. Importantly, these findings extend beyond conventional qualitative assessment of moving intravascular bubbles and suggest that quantitative ultrasound may provide objective biomarkers of severe decompression-related gas burden, including tissue-associated gas accumulation. By leveraging a naturally occurring model of severe systemic gas embolism, this work provides a framework for studying decompression-related ultrasound signatures that are difficult to investigate in human DCS cohorts. This work also has implications for sea turtle conservation efforts. With large numbers of bycaught sea turtles, better understanding gas emboli dynamics and developing tools to better estimate disease severity can assist mitigation efforts to protect these endangered species.

## Acknowledgements

This study was partially supported by the Divers Alert Network (DAN) research foundation through grants DAN-UNC-1 and DAN-UNC-2. The funder had no role in study design, data collection and analysis, decision to publish, or preparation of the manuscript.

## REFERENCES

1. Mitchell SJ. Decompression illness: a comprehensive overview. Diving Hyperb Med. 2024;54(1Suppl):1–53. doi:10.28920/DHM54.1.SUPPL.1-53

2. Mitchell SJ, Bennett MH, Moon RE. Decompression Sickness and Arterial Gas Embolism. N Engl J Med. 2022;386(13):1254–1264.

3. Le DQ, Hoang AH, Azarang A, et al. An open-source framework for synthetic post-dive Doppler ultrasound audio generation. PLoS One. 2023;18(4 April). doi:10.1371/JOURNAL.PONE.0284922

4. Doolette DJ, Murphy FG. Within-diver variability in venous gas emboli (VGE) following repeated dives. Diving Hyperb Med. 2023;53(4):333. doi:10.28920/DHM53.4.333-339

5. Fahlman A. Allometric scaling of decompression sickness risk in terrestrial mammals; cardiac output explains risk of decompression sickness. Scientific Reports 2017 7:1. 2017;7(1):40918-. doi:10.1038/srep40918

6. Papadopoulou V, Germonpré P, Cosgrove D, et al. Variability in circulating gas emboli after a same scuba diving exposure. European Journal of Applied Physiology 2018 118:6. 2018;118(6):1255–1264. doi:10.1007/S00421-018-3854-7

7. Howle LE, Weber PW, Hada EA, Vann RD, Denoble PJ. The probability and severity of decompression sickness. PLoS One. 2017;12(3):e0172665. doi:10.1371/JOURNAL.PONE.0172665

8. Currens JB, Doolette DJ, Murphy FG. Venous gas emboli (VGE) in 2-D echocardiographic images following movement: grading and association with cumulative incidence of decompression sickness. Diving Hyperb Med. 2025;55(1):44–50. doi:10.28920/DHM55.1.44-50

9. Le DQ, Dayton PA, Tillmans F, et al. Ultrasound in decompression research: fundamentals, considerations, and future technologies. Undersea Hyperb Med. 2021;48(1):59–72. doi:10.22462/01.03.2021.8

10. Garcıá-Párraga D, Lorenzo T, Wang T, et al. Deciphering function of the pulmonary arterial sphincters in loggerhead sea turtles (Caretta caretta). J Exp Biol. 2018;221(Pt 23). doi:10.1242/JEB.179820

11. Costello LM, Garcıá-Párraga D, Crespo-Picazo JL, Codd JR, Shiels HA, Joyce W. Absence of atrial smooth muscle in the heart of the loggerhead sea turtle (Caretta caretta): a re-evaluation of its role in diving physiology. Journal of Experimental Biology. 2022;225(20). doi:10.1242/JEB.244864/278473

12. García-Párraga D, Crespo-Picazo JL, De Quirós YB, et al. Decompression sickness (’the bends’) in sea turtles. Dis Aquat Organ. 2014;111(3):191–205. doi:10.3354/DAO02790

13. Sea Turtle | WWF. Accessed July 5, 2026. https://www.worldwildlife.org/species/sea-turtle/

14. Parga ML, Crespo-Picazo JL, Monteiro D, et al. On-board study of gas embolism in marine turtles caught in bottom trawl fisheries in the Atlantic Ocean. Scientific Reports 2020 10:1. 2020;10(1):5561-. doi:10.1038/s41598-020-62355-7

15. Lucchetti A, Sala A. An overview of loggerhead sea turtle (Caretta caretta) bycatch and technical mitigation measures in the Mediterranean Sea. Reviews in Fish Biology and Fisheries 2009 20:2. 2009;20(2):141–161. doi:10.1007/S11160-009-9126-1

16. Fahlman A, Crespo-Picazo JL, Sterba-Boatwright B, Stacy BA, Garcia-Parraga D. Defining risk variables causing gas embolism in loggerhead sea turtles (Caretta caretta) caught in trawls and gillnets. Scientific Reports 2017 7:1. 2017;7(1):2739-. doi:10.1038/s41598-017-02819-5

17. Franchini D, Valastro C, Ciccarelli S, et al. Analysis of risk factors associated with gas embolism and evaluation of predictors of mortality in 482 loggerhead sea turtles. Scientific Reports 2021 11:1. 2021;11(1):22693-. doi:10.1038/s41598-021-02017-4

18. Valastro C, Franchini D, Ciccarelli S, et al. Comparative Diagnostic Efficacy of Ultrasonography and Radiography for Gas Embolism in Loggerhead (Caretta caretta) Turtles. Animals. 2024;14(24):3623. doi:10.3390/ANI14243623/S1

19. Azarang A. User friendly Graphical User Interface for Ultrasound Scan Processing. GitHub. Accessed July 5, 2026. https://github.com/ArianAzg/User-friendly-Graphical-User-Interface-for-Ultrasound-Scan-Processing

20. Haralick RM, Dinstein I, Shanmugam K. Textural Features for Image Classification. IEEE Trans Syst Man Cybern. 1973;SMC-3(6):610–621. doi:10.1109/TSMC.1973.4309314

21. Gómez W, Pereira WCA, Infantosi AFC. Analysis of co-occurrence texture statistics as a function of gray-level quantization for classifying breast ultrasound. IEEE Trans Med Imaging. 2012;31(10):1889–1899. doi:10.1109/TMI.2012.2206398

22. Wang X, Sheng L. Correlations between B-mode ultrasound image texture features and tissue temperatures in hyperthermia. PLoS One. 2022;17(10):e0266446. doi:10.1371/JOURNAL.PONE.0266446

23. Brusasco C, Santori G, Tavazzi G, et al. Second-order grey-scale texture analysis of pleural ultrasound images to differentiate acute respiratory distress syndrome and cardiogenic pulmonary edema. Journal of Clinical Monitoring and Computing 2020 36:1. 2020;36(1):131–140. doi:10.1007/S10877-020-00629-1

24. Alvarenga A V., Teixeira CA, Ruano MG, Pereira WCA. Influence of temperature variations on the entropy and correlation of the Grey-Level Co-occurrence Matrix from B-Mode images. Ultrasonics. 2010;50(2):290–293. doi:10.1016/J.ULTRAS.2009.09.002

25. Currens JB, Moon RE, Makowski MS, et al. Ultrasound detection of lymphatic bubbles in a porcine dive model. J Appl Physiol (1985). 2025;139(2):365–375. doi:10.1152/JAPPLPHYSIOL.00171.2025

26. Papadopoulou V, Eckersley RJ, Balestra C, Karapantsios TD, Tang MX. A critical review of physiological bubble formation in hyperbaric decompression. Adv Colloid Interface Sci. 2013;191-192:22–30. doi:10.1016/J.CIS.2013.02.002

27. Bernaldo de Quirós Y, Møllerløkken A, Havnes MB, Brubakk AO, González-Díaz O, Fernández A. Bubbles quantified in vivo by ultrasound relates to amount of gas detected post-mortem in rabbits decompressed from high pressure. Front Physiol. 2016;7(JUL):207152. doi:10.3389/FPHYS.2016.00310/TEXT

28. Wyneken J. The Anatomy of Sea Turtles. NOAA Technical Memorandum NMFS-SEFSC-470. Preprint posted online December 2001:1–172. Accessed June 24, 2026. https://www.researchgate.net/publication/265924061_The_Anatomy_of_Sea_Turtles_The_Anatomy_of_Sea_Turtles

29. García-Párraga D, Valente ALS, Stacy BA, Wyneken J. Cardiovascular System. In: Charles A. Manire, Terry M. Norton, Brian A. Stacy, Craig A. Harms, Charles J. Innis, eds. Sea Turtle Health and Rehabilitation. 1st ed. 2017:295–320.

30. Stanziola A, Toulemonde M, Yildiz YO, Eckersley RJ, Tang MX. Ultrasound Imaging with Microbubbles. IEEE Signal Processing. Published online March 2016:111–117.

