## Supplemental Figures and Tables for "Quantitative Ultrasound Analysis of Gas Embolic Disease in Bycaught Sea Turtles"

Katherine Eltz, B.S.[1,2], Jose-Luis Crespo, DVM, PhD[3], Arian Azarang, PhD[1,2], Emma Pla-González, DVM[3], Daniel García-Párraga, DVM, PhD, Dipl. ECZM (ZHM); Dipl. ECAAH (N-P) [2,3], Andreas Fahlman, PhD[3,4], Virginie Papadopoulou, PhD[1,2]

Item 1

| Grade | Eftedal-Brubakk Scale |
| --- | --- |
| 0 | No bubbles visible |
| 1 | Occasional bubbles |
| 2 | At least 1 bubble every 4 heartbeats |
| 3 | At least 1 bubble every heartbeat |
| 4 | At least 1 bubble at every $cm^2$ in every view |
| 5 | Whiteout - no single bubble discrimination |

**Table S1:** The Eftedal- Brubakk venous gas emboli (VGE) grading scale used for human echocardiography assessment post dive. Reproduced from [29].

Item 2

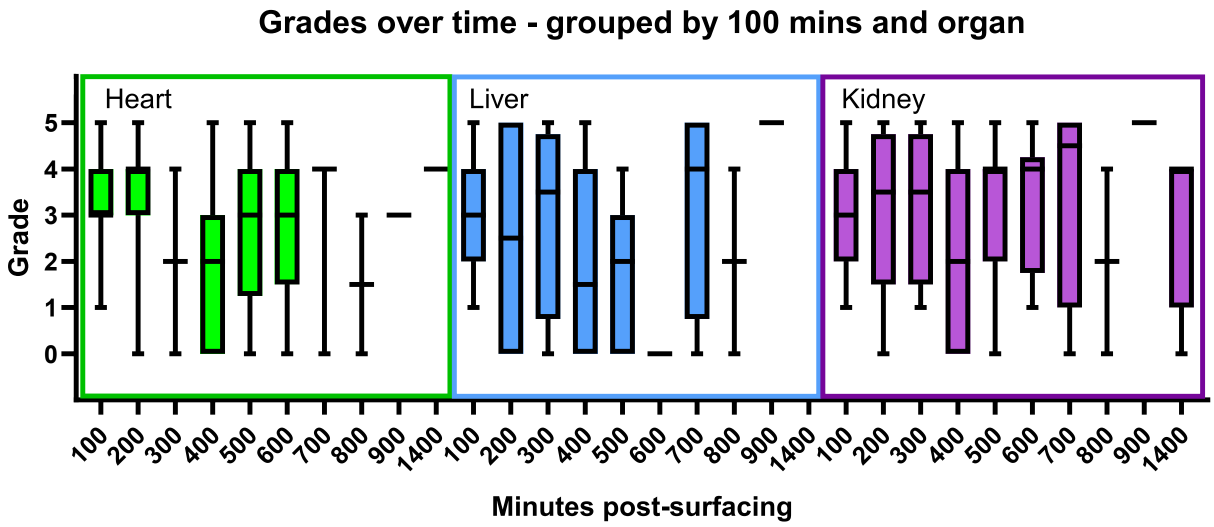


**Figure S1: All recorded bubble grades in the shore and boat groups, separated by organ.** The kidney and liver retained higher bubble grades at later time points, while grade 5 bubbles were no longer observed in the heart at earlier time points than in the other organs.

Item 3

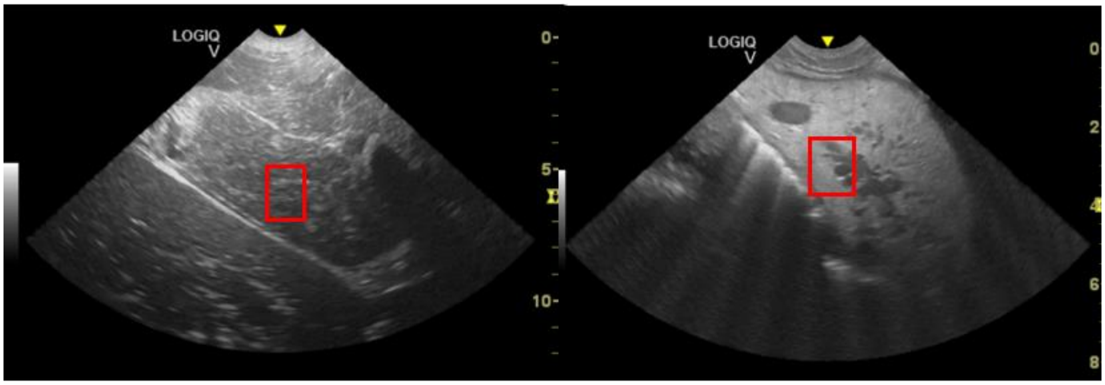


**Figure S2: Example scans of a liver (A) and a kidney (B).** The red squares indicate the region of interest used for texture analysis.

Item 4

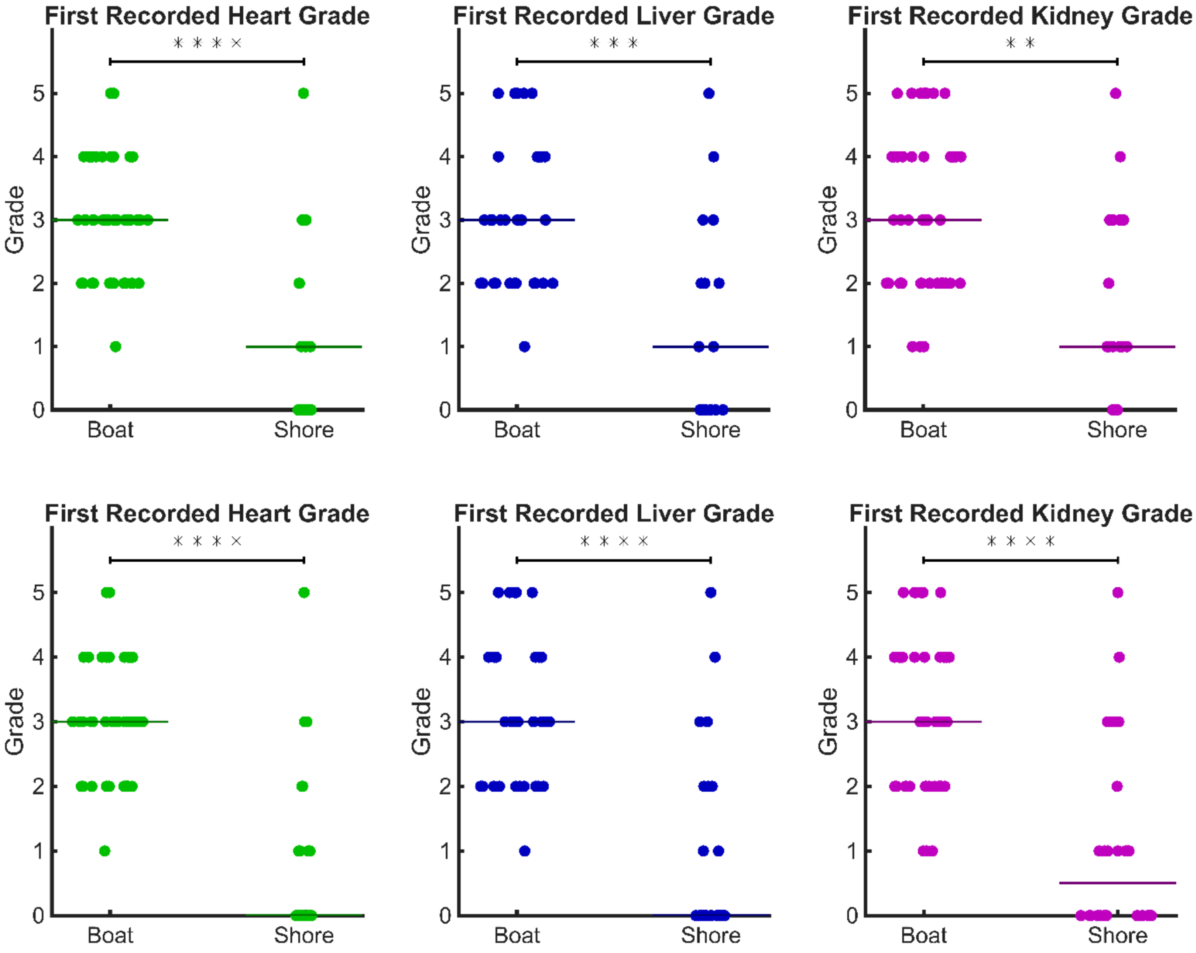


**Figure S3: First recorded bubble grades in both groups, separated by organ. A)** Data from all available trawler-captured turtles with available surface interval. **B)** Data from the subset of turtles with a recorded surface interval, regardless of capture method, including trawling net, trammel net, and stranding.

Item 5

(A) Liver

| Parameters | β0 Intercept | β1 Contrast | β2 Correlation | β3 Energy | β4 Homogeneity |
|---|---|---|---|---|---|
| β0 | 1.00 | | | | |
| β1 | -0.93 | 1.00 | | | |
| β2 | -0.89 | 0.88 | 1.00 | | |
| β3 | -0.79 | 0.85 | 0.80 | 1.00 | |
| β4 | -0.68 | 0.42 | 0.37 | 0.17 | 1.00 |

(B) Kidney

| Parameters | β0 Intercept | β1 Contrast | β2 Correlation | β3 Energy | β4 Homogeneity |
|---|---|---|---|---|---|
| β0 | 1.00 | | | | |
| β1 | -0.74 | 1.00 | | | |
| β2 | -0.70 | 0.52 | 1.00 | | |
| β3 | -0.73 | 0.79 | 0.71 | 1.00 | |
| β4 | -0.73 | 0.29 | 0.14 | 0.16 | 1.00 |

**Table S2**: Parameter covariance matrices for liver (A) and kidney (B) for the multivariate logistic regression model using texture features to predict scan location (shore vs boat).

Item 6

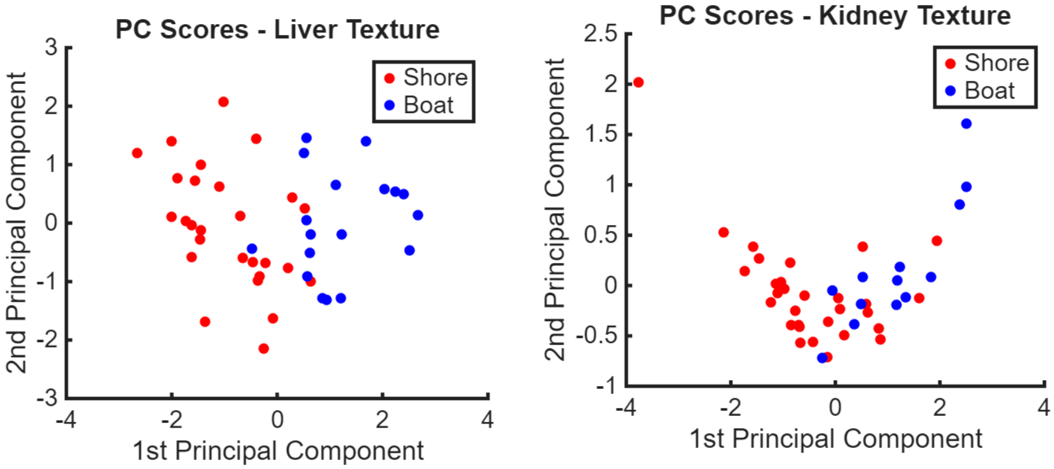


**Figure S5**: Principal component analysis of texture features demonstrating separation between shore and boat scan locations, with scan location used as a proxy for elapsed time.
